# Smoking and blood DNA methylation: novel associations, replication of previous findings and assessment of reversibility

**DOI:** 10.1101/660878

**Authors:** Pierre-Antoine Dugué, Chol-Hee Jung, JiHoon E Joo, Xiaochuan Wang, Ee Ming Wong, Enes Makalic, Daniel F Schmidt, Laura Baglietto, Gianluca Severi, Melissa C Southey, Dallas R English, Graham G Giles, Roger L Milne

## Abstract

We conducted a genome-wide association study of blood DNA methylation and smoking, attempted replication of previously discovered associations, and assessed the reversibility of smoking-associated methylation changes. DNA methylation was measured in baseline peripheral blood samples for 5,044 participants in the Melbourne Collaborative Cohort Study. For 1,032 participants, these measures were repeated using blood samples collected at follow-up, a median of 11 years later. A cross-sectional analysis of the association between smoking and DNA methylation and a longitudinal analysis of changes in smoking status and changes in DNA methylation were conducted. We used our cross-sectional analysis to replicate previously reported associations for current (N=3,327) and former (N=172) smoking. A comprehensive smoking index accounting for the bioactivity of smoking and several aspects of smoking history was constructed to assess the reversibility of smoking-induced methylation changes. We identified 4,496 cross-sectional associations at P<10^−7^, including 3,296 that were novel. We replicated the majority (90%) of previously reported associations for current and former smokers. In our data, we observed for former smokers a substantial degree of return to the methylation levels of never smokers, compared with current smokers (median: 74%, IQR=63-86%). Consistent with this, we found wide-ranging estimates for the half-life parameter of the comprehensive smoking index. Longitudinal analyses identified 368 sites at which methylation changed upon smoking cessation. Our study provides evidence of many novel associations between smoking and DNA methylation at CpGs across the genome, replicates the vast majority of previously reported associations, and indicates wide-ranging reversibility rates for smoking-induced methylation changes.

## INTRODUCTION

Several studies have examined the association between exposure to tobacco smoke and DNA methylation levels in blood.^**1–13**^ A systematic review identified methylation at 1,460 CpG sites to be associated with smoking,^**14**^ and a recent large-scale study identified 2,623 CpGs with P<10^−7^.^**12**^ These associations were identified comparing current with never smokers, and not all were replicated using independent data. Additionally, there is substantial variability by study in the strength of associations, which may be due to characteristics of the cohorts such as age or ethnicity, or methodological issues such as the variables used for adjustment in statistical models or the pipeline used for normalisation of the DNA methylation data.

Most of these studies also reported differences in methylation for former smokers compared with never and current smokers, indicating a degree of reversibility of smoking-associated methylation changes. Few studies have examined reversibility patterns beyond assessing the effect of time since quitting.^**5, 10, 12**^ Guida and colleagues assessed reversibility in a study based on 745 women and identified two clusters of smoking-associated methylation at CpG sites according to whether methylation reverted back to the level of never smokers within 35 years of quitting.^**5**^ The assessment of reversibility made by Joehanes and colleagues was based on 2,374 participants and concluded that for the majority of the 2,568 CpGs they examined (those with FDR-adjusted P<0.05 in the comparison of former vs. never smokers) methylation levels returned to those of never smokers within five years of smoking cessation, and for only 36 CpGs did they observe no tendency of a return to the methylation levels of never smokers 30 years after they had quit.^**12**^ Consistent findings were reported by Wilson and colleagues, who made use of repeated methylation measures taken seven years apart to identify methylation at CpG sites that varied longitudinally with changes in smoking status.^**10**^ They also observed differential methylation in former smokers who had quit more than 40 years before methylation measurement, compared to never smokers. Assessing what smoking-associated methylation changes are transient or long-lasting may have important implications for biological understanding and clinical practice.^**15**^

The bioactivity of exposure to smoking can be modelled as a function of the smoking history of an individual, including the number of cigarettes smoked, the age at starting smoking, and the duration of smoking. The resulting comprehensive smoking index (CSI) was shown to substantially improve the prediction of smoking-related disease compared with simpler smoking assessment models.^**16–18**^ A prominent feature of the CSI is that it includes a parameter for biological half-life, representing the rate at which the activity of smoking compounds declines, and is therefore the parameter of interest when assessing reversibility.

In this study, we aimed to: i) conduct a genome-wide association study of DNA methylation and exposure to tobacco smoking measured using traditional smoking assessment and CSI,^**16**^ the latter allowing a better assessment of the methylation reversibility pattern; ii) replicate previously reported associations, including associations observed in former smokers or by time since quitting; iii) assess the association between changes in DNA methylation and changes in smoking using repeated measures taken a median of 11 years apart.

## MATERIALS AND METHODS

### Study participants

Between 1990 and 1994 (baseline), 41,513 participants were recruited to the Melbourne Collaborative Cohort study (MCCS). The majority (99%) were aged 40 to 69 years and 41% were men. Southern European migrants were oversampled to extend the range of lifestyle factors and genetic variation^**25**^ Participants were contacted again between 2003 and 2007 (follow-up). Blood samples were taken at baseline and follow-up from 99% and 64% of participants, respectively. Baseline samples were stored as dried blood spots on Guthrie cards for the majority (73%), as mononuclear cell samples for 25% and as buffy coat samples for 2% of the participants. Follow-up samples were stored as buffy coat samples and dried blood spots on Guthrie cards. All participants provided written informed consent and the study protocols were approved by the Cancer Council Victoria’s Human Research Ethics Committee.

The present study sample comprised MCCS participants selected for inclusion in one of seven previously conducted nested-case control studies of DNA methylation of colorectal, gastric, kidney, lung, and prostate cancer, B-cell lymphoma, and urothelial cell carcinoma (UCC).^**26–31**^ Controls were matched to incident cases of prostate, colorectal, gastric, lung or kidney cancer, UCC or mature B-cell neoplasms on sex, year of birth, country of birth, baseline sample type and smoking status (the latter for the lung cancer study only). For the cross-sectional analyses, we excluded participants whose blood sample was taken at follow-up (303 samples from the UCC study) because their questionnaire data and storage time were different. We also excluded cases from the lung and UCC studies to avoid bias due to the strong association between smoking and these cancers.^**24**^ Methylation data for baseline blood samples (baseline study) were available from a total of 2,777 controls and 2,267 cases after quality control and exclusions. Additionally, methylation measures (Guthrie cards) were repeated at follow-up (2004-2007) for a subset of 1,100 of the controls who also had their baseline sample collected on a Guthrie card, of which 1,088 were available after quality control.

Description of the smoking variables is presented in **Table 1**. Participants with missing data for smoking variables were excluded from the analysis, as were those who had never smoked cigarettes but had smoked cigars or pipes. Missing data for confounders (<1% for age, sex, ethnicity, BMI or alcohol drinking) were imputed using the median or mode of the distribution for continuous and categorical variables, respectively.

**Table 1.** Characteristics of study participants from the Melbourne Collaborative Cohort Study (MCCS) at baseline (1990-1994) and follow-up (2003-2007)

|  | Cross-sectional analysis<br>(N=5,044) | Longitudinal analysis<br>(N=1,024) |  |
| --- | --- | --- | --- |
|  | Baseline data | Baseline data | Wave 2 data |
| Age in years, median, interquartile range [IQR] | 60.7 [53.9-65.4] | 58.5 [51.1-64.1] | 69.8 [62.7-75.5] |
| Sex, male | 3,408 (68%) |  | 701 (68%) |
| Country of birth |  |  |  |
| AU/NZ/Other | 3,411 (68%) |  | 799 (78%) |
| Greece | 382 (8%) |  | 36 (4%) |
| Italy | 714 (14%) |  | 75 (7%) |
| UK | 537 (11%) |  | 114 (11%) |
| BMI (kg/m<sup>2</sup>), median [range] | 26.9 [24.5-29.5] | 26.3 [24.1-29.0] | 26.7 [24.2-29.3] |
| Alcohol intake (g/day), median [IQR] | 4.3 [0.0-18.7] | 4.3 [0.0-18.6] | 7.9 [0.3-22.7] |
| Smoking status |  |  |  |
| Never | 2,379 (47%) |  |  |
| Former ≥15 years ago | 1059 (21%) |  |  |
| Former <15 years ago | 951 (19%) |  |  |
| Current <20 cig/day | 269 (5%) |  |  |
| Current ≥20 cig/day | 386 (8%) |  |  |
| Smoking status at baseline and follow-up |  |  |  |
| Never-Never |  |  | 518 (51%) |
| Former-Former |  |  | 400 (39%) |
| Current-Former |  |  | 50 (5%) |
| Current-Current |  |  | 56 (5%) |

Methods relating to DNA extraction, and DNA methylation processing and quality are presented in Supplementary Material.

### Previously reported associations

We identified previous studies using the keywords (“smoking” and “blood” and “methylation”), which returned 416 articles in PubMed (31 July 2018). We retained from this search six studies having conducted an EWAS of smoking and blood DNA methylation.^**2, 5, 9, 10, 12, 32, 33**^ Other studies were identified but not selected due to small sample size (N<200), or not adjusting for potential confounders of the association.^**1, 3, 4, 6–8, 11, 34–38**^ The six studies retained identified 3,327 associations with a P-value less than 10^−7^, 2500 (75%) in one study only, 438 (13%) in two, and 389 (12%) in three or more studies. Of the six studies, four also reported differentially methylated CpGs for former compared with never smokers,^**5, 9, 12, 32**^ identifying 172 associations, including 146 in only one study.

### Comprehensive smoking index (CSI)

We constructed a CSI following the recommendations of Leffondré and colleagues.^**16**^ We observed better model fits (data not shown) when using the log-transformed version of the CSI: ln(CSI)+1, referred to as simply ‘CSI’, and we assumed no lag-time between exposure to smoking and changes in DNA methylation.^**16**^ The CSI was defined in our study as:

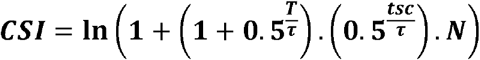

where *T* is duration of smoking in years, *tsc* the time since smoking cessation in years, *N* the average number of cigarettes smoked per day and *τ* the half-life parameter. We estimated *τ* from the data as follows: (i) by visual inspection of CSI values obtained for various *τ* values (***Figure 1***), we concluded that for smaller values of *τ*, the CSI was both sensitive and more consistent with assumed biological activity by smoking history; (ii) for a CpG of interest, we fitted the same model for every CSI with *τ* value within the grid: {0.001; 0.005; 0.01; 0.025; 0.05; 0.1 to 1 by increment of 0.1; 1 to 10 by increment of 0.25; 10 to 30 by increment of 1; and 30 to 100 by increment of 10}; (iii) the estimated τ that maximised model fit,^**16**^ based on the restricted maximum likelihood from a linear mixed model (see following section).

**Figure 1.**
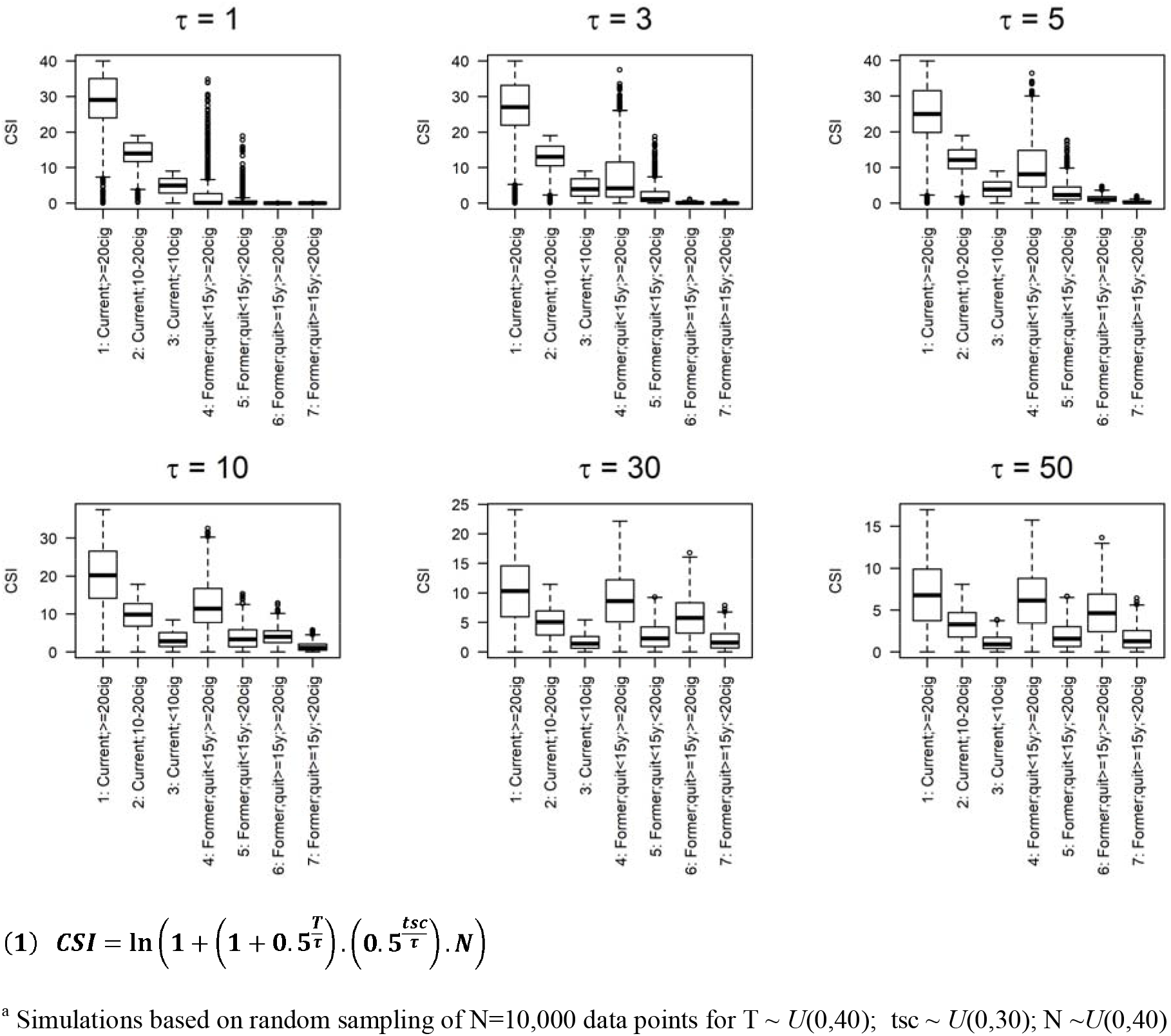
Relationship^a^ between smoking status and comprehensive smoking indices (equation (1)) for various values of half-life parameter τ.

### Genome-wide association study of DNA methylation (EWAS)

We assessed cross-sectional associations (baseline data) for methylation at each individual CpG by regressing DNA methylation M-values on smoking status using linear mixed-effects regression models, using the function *lmer* from the R package *lme4*.^**39**^ Models were adjusted by fitting fixed effects for baseline values of age (continuous), country of birth (Australia/New-Zealand, Italy, Greece, United Kingdom/Malta), sex, alcohol drinking in the previous week (continuous, in grams/day), BMI (≤25 kg/m2, >25 to ≤30, >30 to ≤35, >35), sample type (peripheral blood mononuclear cells, dried blood spots, buffy coats) and estimated white blood cell composition (percentage of CD4+ T cells, CD8+ T cells, B cells, NK cells, monocytes and granulocytes, estimated using the Houseman algorithm^**21, 22**^), and random effects for study, plate, and chip. Heterogeneity in the association between smoking and methylation by age (continuous), sex, alcohol intake in the previous week (continuous), BMI (continuous) and future case status was assessed using likelihood ratio tests for interaction.

We estimated *τ* for the 3,327 CpGs previously reported to be associated with smoking. These findings are summarised in **Table 4** and **Figure 1**. We assumed that the median, and the 25^th^ and 75^th^ percentile of the distribution of τ were the values most likely to detect novel associations between smoking and DNA methylation. We thus ran cross-sectional EWAS analyses for: i) current compared with never smoking, ii) former compared with never smoking; and iii) CSI (continuous variable) with τ=1.5, τ=2.75, and τ=5.25. Given the substantial correlation between these tests, we did not correct further for multiple testing and used a threshold of P<10^−7^ to identify associations for any of these EWAS.^**40–42**^ The false discovery rate (FDR-adjusted P<0.05) was used to identify suggestive associations.^**12, 32**^

For all associations with P<10^−7^ in our cross-sectional EWAS we estimated the half-life τ that provided the best model fit for the CSI, as described previously. We also calculated a ‘reversibility coefficient’, expressed as a percentage and defined as the regression coefficient comparing ‘*former*’ to ‘*current*’ smokers divided by the coefficient comparing ‘*never*’ to ‘*current*’ smokers, as done previously.^**20**^

### Longitudinal analysis

Linear mixed effects regression models were used to assess the relationship between change in smoking status and change in methylation for individual differentially methylated CpGs in our crosssectional EWAS (P<10^−7^). In a first model, we used the following longitudinal smoking patterns: current (at baseline)-current (at follow-up), current-former, former-former, and never-never. Study was included as a random effect and the following variables were included as fixed effects: sex, country of birth (four categories), baseline age (continuous), baseline alcohol intake (continuous), baseline BMI (continuous), baseline cell composition (as defined previously), change in age, BMI and alcohol intake (all continuous), the difference between baseline and follow-up composition for each cell type (continuous), baseline smoking (expressed using a CSI with *τ*=1.5 because it identified the greatest number of associations in the cross-sectional EWAS) and the baseline methylation M-value of the CpG. As adjustment for baseline methylation in analyses of change in methylation may lead to bias in some circumstances,^**43**^ we conducted a sensitivity analysis using models without adjustment for baseline M-value. We also carried the analysis not adjusting for baseline smoking status.

All statistical analyses were performed using the statistical software R (version 3.4.4).

## RESULTS

Altogether, 5,044 participants in the Melbourne Collaborative Cohort Study (MCCS) were included in the cross-sectional analysis; at baseline, their median age was 60.7 years (IQR: 53.9-65.4), 3,408 (68%) were males, and 655 (68%) were current, 2,010 (40%) former, and 2,379 (47%) never smokers (**Table 1**). Participants in the longitudinal analysis were younger (median age at baseline: 58.5 years) and generally had healthier lifestyle than other participants included in the cross-sectional analysis.

### Genome-wide association study of DNA methylation

#### Comparison of current, former and never smokers

At P<10^−7^, we observed 1,851 differentially methylated CpG sites between current and never smokers, and 156 differentially methylated CpGs between former and never smokers, with 140 overlapping CpGs and 16 found in former smokers only. In total, 917 of the 1,851 CpGs (50%) associated with current smoking had not been reported in previous studies at P<10^−7^ (**Supplementary Table 1**); 1,124 (61%) showed some methylation differences (P<0.05 and same direction of coefficient) in former smokers. Reversibility coefficients indicated that for former smokers, there was a substantial degree of return to methylation levels of never smokers (median: 74%, IQR=63% to 86%).

#### Comprehensive smoking indices (CSI)

We first considered plausible values of half-life parameter τ of the CSI based on 3,327 differentially methylated CpGs identified in six previous studies at P<10^−7^ (**Supplementary Table 2**). Estimated *τ* values were wide-ranging: median: 2.25, IQR: 1 to 5.25 and 3,038 (91%) CpGs had P<0.05. To further refine the potential for these values to identify new associations, we considered only the 1,277 CpGs for which the previously reported association was replicated in our sample (with the estimated *τ*) at P<10^−7^. For these, the median and 25^th^ and 75^th^ percentile values were 2.75, 1.5 and 5.25 respectively. These values were consistent with the simulated values presented in **Figure 1**. We thus conducted methylome-wide association studies for each of these three values and identified 3,497 (*τ*=2.75), 4,022 (*τ*=1.5) and 2,433 (*τ*=5.25), respectively, at P<10^−7^. From these analyses, 4,496 associations were identified and DNA methylation at these CpGs was classified as smoking-associated in subsequent analyses, including 1,775 overlapping with associations identified using the current and former smoking variables. Of these, 3,296 (73%) had not been reported at P<10^−7^ in previous studies.

#### Interaction analyses

Using the Bonferroni correction for multiple testing (P=0.05/4,496=1.1×10^−5^) and the CSI with *τ*=1.5, we observed a weaker association for DNA methylation in women at a CpG not annotated to a gene, and a weaker association for participants with higher BMI at five CpGs, including two in *AHRR* (**Supplementary Table 3**). No significant interaction with smoking status was observed at this significance threshold by age, alcohol consumption, or future case status.

#### Replication of previously reported associations

We examined the replication in the MCCS of 3,327 associations between current smoking and whole-blood DNA methylation previously reported in any of the six studies considered. We replicated, with coefficients consistent in direction, 2,795 (84%) at P<0.05 and 934 (28%) at P<10^−7^ using the current vs. never comparison. These numbers were 2,946 (89%) and 1,200 (36%), respectively, when considering any of the CSIs with τ=1.5, τ=2.75 or τ=5.25 (**Table 2, Supplementary Table 2**). Of the 2,500 associations that had been reported in one study only, we replicated 1,983 (79%) at P<0.05 using the current smoking variable; and 97% of associations that had been reported in two or more studies (**Table 2**).

**Table 2.**
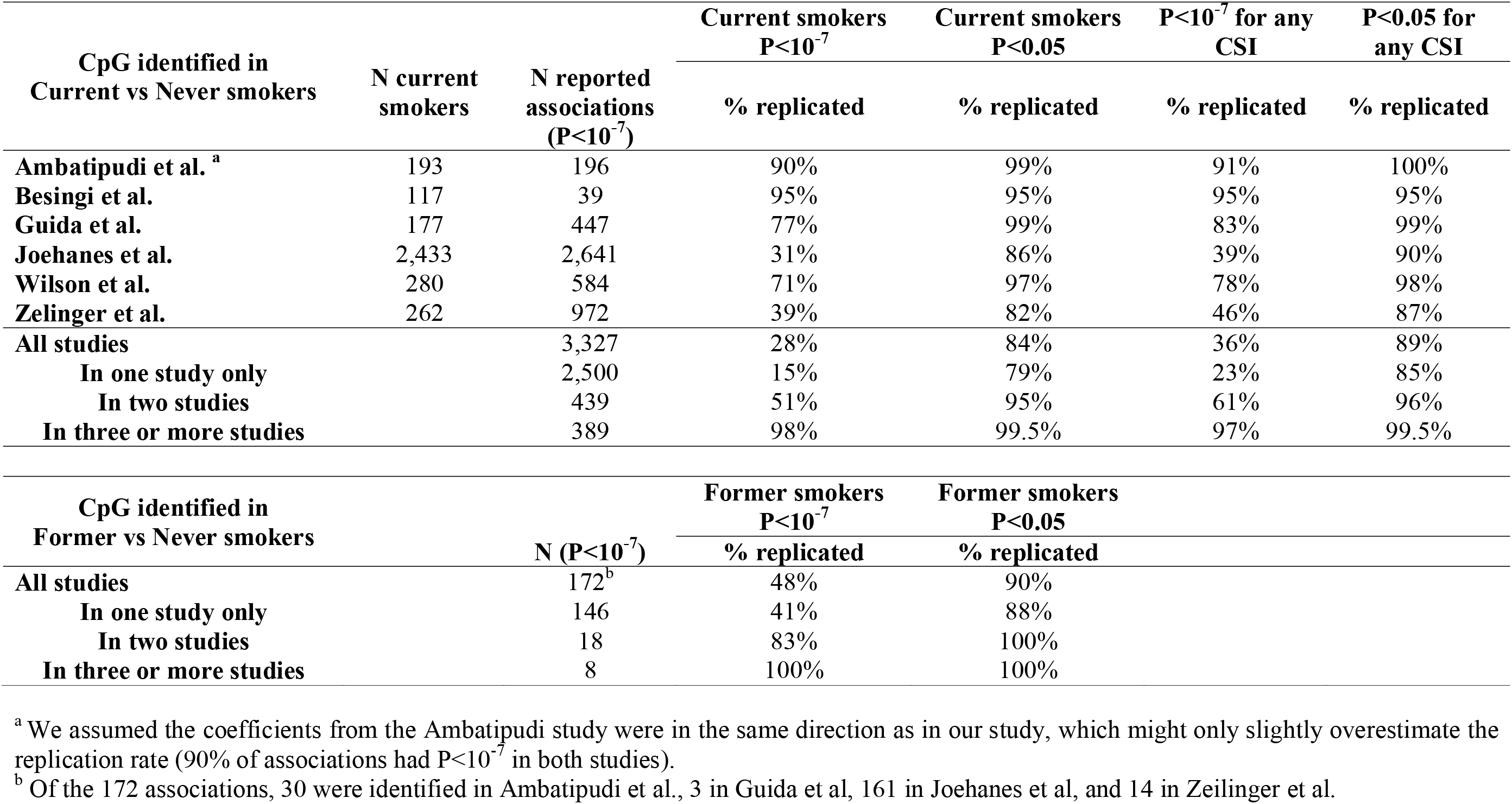
Replication of previously reported associations (P<10^−7^ in six large studies) using the ‘Current vs Never’ smoker comparison, and for 3 comprehensive smoking indices (τ=1.5, τ=2.75 or τ=5.25)

| CpG identified in<br>Current vs Never smokers | N current<br>smokers | N reported<br>associations<br>($P < 10^{-7}$) | Current smokers<br>$P < 10^{-7}$ | Current smokers<br>$P < 0.05$ | $P < 10^{-7}$ for any<br>CSI | $P < 0.05$ for<br>any CSI |
| --- | --- | --- | --- | --- | --- | --- |
|  |  |  | % replicated | % replicated | % replicated | % replicated |
| Ambatipudi et al. <sup>a</sup> | 193 | 196 | 90% | 99% | 91% | 100% |
| Besingi et al. | 117 | 39 | 95% | 95% | 95% | 95% |
| Guida et al. | 177 | 447 | 77% | 99% | 83% | 99% |
| Joehanes et al. | 2,433 | 2,641 | 31% | 86% | 39% | 90% |
| Wilson et al. | 280 | 584 | 71% | 97% | 78% | 98% |
| Zelinger et al. | 262 | 972 | 39% | 82% | 46% | 87% |
| All studies |  | 3,327 | 28% | 84% | 36% | 89% |
| In one study only |  | 2,500 | 15% | 79% | 23% | 85% |
| In two studies |  | 439 | 51% | 95% | 61% | 96% |
| In three or more studies |  | 389 | 98% | 99.5% | 97% | 99.5% |

| CpG identified in<br>Former vs Never smokers | N ($P < 10^{-7}$) | Former smokers<br>$P < 10^{-7}$ | Former smokers<br>$P < 0.05$ |
| --- | --- | --- | --- |
|  |  | % replicated | % replicated |
| All studies | 172<sup>b</sup> | 48% | 90% |
| In one study only | 146 | 41% | 88% |
| In two studies | 18 | 83% | 100% |
| In three or more studies | 8 | 100% | 100% |

<sup>a</sup> We assumed the coefficients from the Ambatipudi study were in the same direction as in our study, which might only slightly overestimate the replication rate (90% of associations had $P < 10^{-7}$ in both studies).
<sup>b</sup> Of the 172 associations, 30 were identified in Ambatipudi et al., 3 in Guida et al, 161 in Joehanes et al, and 14 in Zeilinger et al.

We then examined the replication of associations identified for former compared with never smoking previously reported in any of four large studies. Of the 146 associations that had been reported at P<10^−7^ in one study only, we replicated 129 (88%) at P<0.05 and 60 (41%) at P<10^−7^ using the former smoking variable. All associations that had been reported two or more times were replicated at P<0.05 using the MCCS data (**Table 2, Supplementary Table 4**).

#### Replication of our findings

We examined the replication of our findings using the results from Joehanes et al.^**12**^ in which P-values up to 0.019 (FDR-adjusted P<0.05) were presented for the current vs. never smoking association. Of the 3,296 associations that were novel in our study (P<10^−7^), 1,189 (36%) were replicated at P<0.019 with effect estimates in the same direction.

### Reversibility of associations

Estimated τ values for the 4,496 associations were wide-ranging (**Supplementary Table 5**) but 90% were less than 6, with median [IQR] of 1.75 [1.25-3], consistent with **Figure 1** and the 3,327 previously reported associations. The median τ was equal to 2 for CpGs that were differentially methylated in current or former smokers, compared with never smokers. **Figure 2** shows the relationship between estimated values of τ and: i) reversibility coefficients; this analysis showed greater values of τ for CpGs at which methylation levels in former smokers were similar to those of current smokers, and ii) the strength of association observed in current compared with never smokers; this analysis showed slightly greater τ values for most strongly differentially methylated CpGs in the cross-sectional EWAS.

**Figure 2.**
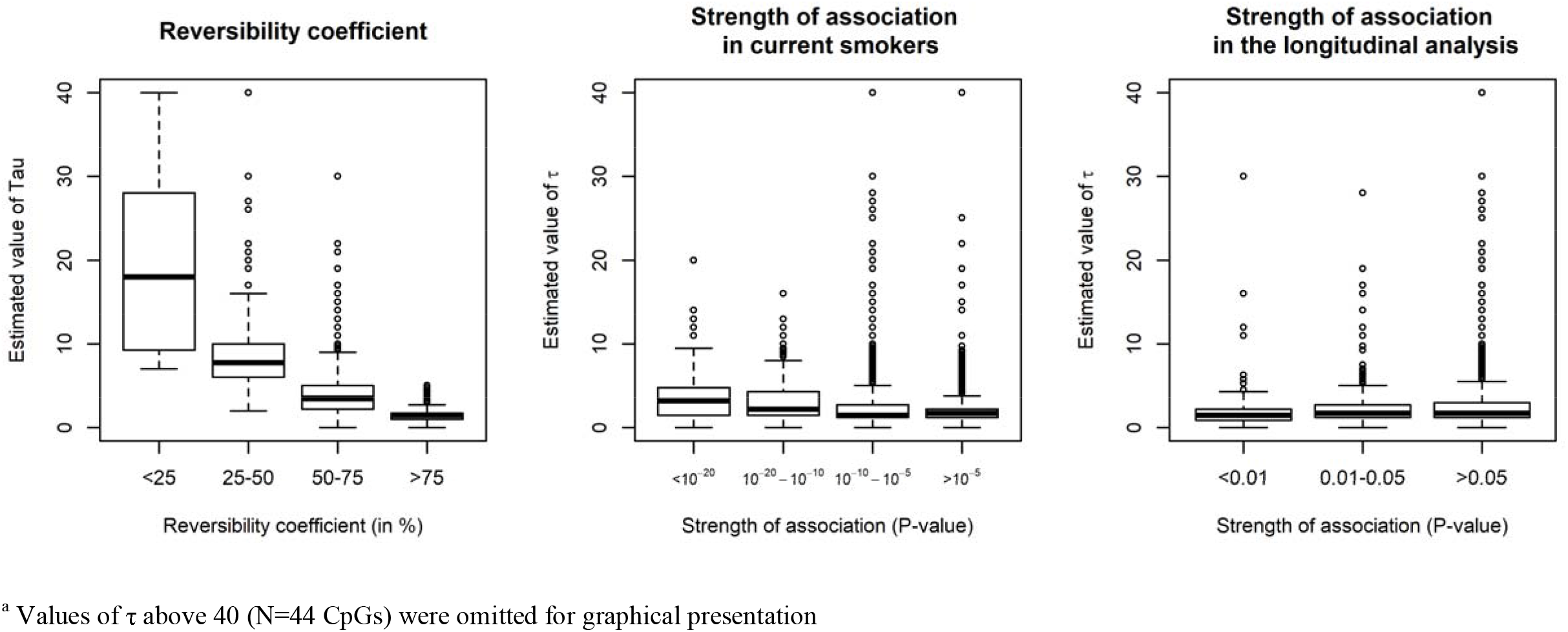
Estimated values of half-life parameter τ according to other features of smoking-associated CpG sites^a^

We then examined the distribution of τ values according to the reversibility patterns observed in three previous studies. First, Guida et al.^**5**^ grouped differentially methylated CpGs into persistent (N=149) or reversible (N=602) clusters. We found weak evidence (Wilcoxon rank-sum test onesided P=0.03) that τ values were greater in the persistent cluster (median τ (IQR): 3.75 [1.75-5.25]) compared with the reversible cluster (2.75 [1.75-5.00]) Second, Joehanes et al.^**12**^ identified 36 CpGs at which methylation levels did not return to never-smoker levels 30 years after smoking cessation: for these CpGs, we found τ values that were greater than for other differentially methylated CpGs (6.25 [3.25-13], one-sided P<0.001). Third, in Wilson et al.^**10**^, 15 CpGs were differentially methylated in participants who had quit smoking for 40 years or more: for these 15 CpGs, we found weak evidence (one-sided P=0.05) of greater τ values (3.75 [2.5-5.25] than for other differentially methylated CpGs.

We further examined the 4,496 cross-sectional associations for longitudinal associations using repeated methylation measures and smoking information collected a median of 11 years apart. After adjustment for baseline smoking status (CSI with τ = 1.5), the results were, comparing with smokers at both time points, 368 differentially methylated CpGs (P<0.05) in participants who had quit from baseline to follow-up, 280 differentially methylated CpGs in former smokers at baseline and 262 in never smokers. The regression coefficients for current-to-former and former-to-former smokers were a median 35% and 90%, respectively, those observed for never smokers. The results without adjustment for baseline M-value were qualitatively similar, albeit identifying fewer longitudinal associations (**Supplementary Table 6a**).

When no adjustment for baseline smoking status was made, compared with participants who were smokers at both time points, 432 CpGs were differentially methylated (P<0.05) in participants who had quit between baseline and follow-up, 1,233 differentially methylated CpGs in former smokers at baseline, and 1,495 in never smokers; regression coefficients for current-to-former and former-to-former smokers were a median 56% and 89%, respectively, that of never smokers (**Supplementary Table 6b**). Using the results with adjustment for baseline smoking status and baseline DNA methylation, we found no evidence that most strongly differentially methylated CpGs in current-to-former compared with current smokers at both time points had lower τ values (**Figure 2**).

## DISCUSSION

Our study identified several thousand novel differentially methylated CpG sites with respect to smoking; 3,296 CpGs with P<10^−7^ that had not been reported at this threshold before were discovered in our cross-sectional EWAS and 1,189 (36%) of these were replicated using the results from a previous large study.^**12**^ The findings using a less conservative significance threshold (FDR) indicate that many more associations exist across the genome, but these would likely be of smaller magnitude, hence possibly less replicable and biologically relevant. This is consistent with the relatively lower replication rate observed for CpGs discovered using the FDR in study by Joehanes et al.,^**12**^ and a simulation study that estimated an optimal multiple testing correction threshold for the HM450 assay to be 2.4×10^−7^.^**19**^

Although the replication of our novel associations may appear relatively low, it should be noted that ‘low-hanging fruit’ were already discovered by previous studies. A testament to the quality and scientific value of our study is the substantial replication we observed for findings of previous studies (~80% for associations reported only once, and 97% for associations reported twice or more). Our literature review might have missed some previously discovered smoking-associated methylation measures at CpGs, but we likely included the majority of them. Additionally, for former smoking, we replicated a substantial proportion (90%) of previously reported associations and identified many novel differentially methylated CpGs.

We assessed associations using a comprehensive smoking index to account for the bioactivity of various smoking exposures relevant to DNA methylation. This modelling strategy has several limitations, including our assumptions that there was no lag-time between smoking exposure and changes in DNA methylation, and that the number of cigarettes smoked contributed equally to methylation changes throughout the lifetime. Another limitation is that because the CSI was log-transformed, the interpretation of the parameter τ was no longer that of a biological half-life, i.e. the time required for a biological substance to reduce to half its initial value.^**16**^ Specific to our study, this means that τ is not interpretable as half the time by which methylation levels of former smokers would return to the level of never smokers. Our values can nevertheless be used to rank CpGs by their rate of reversibility. We also observed (by definition) a clear correspondence between the values of τ and the reversibility coefficients we calculated, suggesting that our analysis provides a more complete picture of how smoking-associated methylation changes vary over time. The main strength of the CSI is that it captures in a single variable several aspects of a smoking history that individually contribute to differential methylation, hence resulting in a more accurate measure of the effects of smoking (illustrated by e.g. >4,000 CpGs identified with τ=1.5, which was substantially more that with the current smoking variable). Finally, the reversibility coefficients calculated in this study were substantially lower than those observed in our previous analysis of alcohol consumption,^**20**^ which suggests that smoking-associated methylation marks might be more frequent but less persistent compared to alcohol-associated methylation changes.

Our longitudinal analysis had less precision due to fewer participants with relatively small variation in smoking status over a decade in this age range, and there was no clear correspondence with reversibility patterns observed from the cross-sectional data. It nevertheless identified many CpGs at which methylation levels returned toward normal in participants who had quit at follow-up compared with those still currently smoking, but these findings need to be replicated. Another limitation of our study is the potential for residual confounding, especially by white blood cell type composition, which is strongly associated with smoking and DNA methylation. Cell composition was estimated with the widely used Houseman algorithm^**21, 22**^ and we did not assess sensitivity to the method used for deriving cell composition.^**23**^ Additionally, we reported in a previous study that many differentially methylated CpGs with respect to alcohol drinking are also associated with smoking, so it may be difficult to tease out the individual effects or joint influences on many of these CpGs across the genome.^**20**^ Finally, we included who later developed cancer, which could give rise to collider bias given the strong association of smoking with cancer risk,^**24**^ but, by assessing effect modification by case-control status, we found no evidence of such bias in our setting.

To conclude, our study provides evidence that several thousand associations between smoking and DNA methylation at CpGs exist across the genome that had not been discovered or replicated before. Smoking-associated methylation changes appeared largely reversible after smoking cessation. We also proposed a way to quantify the reversibility of methylation changes due to smoking by using a comprehensive smoking index that accounts for both the bioactivity of smoking and several aspects of smoking history that are relevant to DNA methylation.

## Acknowledgements and funding

This work was supported by the Australian National Health and Medical Research Council (NHMRC) [grant 1088405]. MCCS cohort recruitment was funded by VicHealth and Cancer Council Victoria. The MCCS was further supported by Australian NHMRC grants 209057, 251553 and 504711 and by infrastructure provided by Cancer Council Victoria. Cases were ascertained through the Victorian Cancer Registry (VCR) and the Australian Cancer Database (Australian Institute of Health and Welfare). The nested case-control methylation studies were supported by the NHMRC grants 1011618, 1026892, 1027505, 1050198, 1043616 and 1074383. M.C.S. is an NHMRC Senior Research Fellow (1155163).

## Notes

#### Summary of Updates

Minor edits.

