## Supplementary methods for "Smoking and blood DNA methylation: novel associations, replication of previous findings and assessment of reversibility"

**DNA methylation data**

***Bisulphite conversion of DNA and Illumina Infinium methylation assay***

Genomic DNA was extracted from mononuclear cells using the QIAamp 96 DNA blood kit (Qiagen), and from Guthrie card samples as previously described **(****Joo *et al.* 2013****).** Genomic DNA was bisulphite converted using the EZ DNA Methylation-Gold kit (Zymo Research, Irvine, CA) following the manufacturer’s instructions. The Illumina Infinium HumanMethylation450K BeadChip (HM450K) array (Illumina, Inc.; San Diego, CA, USA) was used to measure DNA methylation. This array covers 99% of the RefSeq genes and uses a combination of two distinct probe types (Infinium I and II) to detect the methylation status of 485,577 CpGs in the human genome at a single base resolution **(****Bibikova *et al.* 2011****).** Samples in each nested case-control study and the longitudinal study were assayed at non-overlapping time periods. In each study, samples were randomly assigned to chips and processed as per the Illumina protocol. All laboratory work, including DNA extraction, bisulphite conversion and Infinium assaying was performed at the Genetic Epidemiology Laboratory, University of Melbourne.

***Normalization and quality control of methylation data***

The same normalization and quality control procedures were applied to methylation data in all eight studies (case-control and longitudinal). Raw .IDAT files were imported into R using the *minfi* package **(****Aryee *et al.* 2014****).** Illumina’s background correction was applied based on internal control probes. Subset-quantile within-array normalisation (SWAN) was used to correct for the technical variability between the two probe types of the HM450K array **(****Maksimovic *et al.* 2012****)**. The ‘getSex’ function of the *minfi* package **(****Aryee *et al.* 2014****)** was used to predict sex for each sample. Samples for which predicted sex was inconsistent with recorded sex were excluded from further analyses. For each sample, each CpG with a detection *P* value >0.01 was assigned as missing. Samples with missing values for more than 5% of probes were excluded. CpG sites were excluded if they were missing for more than 20% of samples. β-values, which range from zero to one and correspond to the percentage of methylation, were calculated for each CpG using *minfi*. β-values were transformed into M-values using the formula: $M= {log}_{2}(\frac{\beta}{1-\beta})$. **(****Du *et al.* 2010****)**. For the longitudinal analysis (see below), ComBat **(****Johnson *et al.* 2007****)** was applied to remove batch effects. Because ComBat does not accommodate missing values, they were first imputed to the CpG-specific median methylation value and later converted back to missing.
